# PHYSIOLOGICAL CHARACTERISTICS OF BACTERIAL DROPLETS INDICATE A CATASTROPHIC CONSEQUENCE WITH AN INCREASE IN IMPACT VELOCITY

**DOI:** 10.1101/2022.05.28.493826

**Authors:** Vishnu Hariharan, Atish Roy Chowdhury, Srinivas Rao S, Dipshikha Chakravortty, Saptarshi Basu

## Abstract

Droplet impacts on various surfaces play a profound role in different bio-physiological processes and engineering applications. The current study opens a new realm that investigates the plausible effect of impact velocities on bacteria-laden droplets against a solid surface. We unveiled the alarming consequences of *Salmonella* Typhimurium (STM) laden drop, carrying out the *in vitro* and intracellular viability of STM to the impact Weber numbers ranging from 100-750. The specified Weber number range mimics the velocity range occurring during the respiratory processes, especially the airborne dispersion of drops during cough. A thick ring of bacterial deposition was observed in all cases irrespective of impacting velocity and the nutrient content of the bacterial medium. The mechanical properties of the bacterial deposit examined using Atomic Force Microscopy reveals the deformation of bacterial morphology, cushioning effect and adhesion energy to determine the cell-cell interactions. The impact velocity induces the shear stress onto the cell walls of STM, thereby deteriorating the *in vitro* viability. However, we found that even with compromised *in vitro* viability, *Salmonella* retrieved from deposited patterns impacted at higher velocity revealed an increased expression of *phoP* (the response regulator of the PhopQ two-component system) and uninterrupted intracellular proliferation in macrophages. The inability of STM *ΔphoP* growth in nutrient-rich dried droplets to the subjected impact velocities signifies the predominant role of *phoP* in maintaining the virulence of *Salmonella* during desiccation stress. Our findings open a promising avenue for understating the effect of bacteria-laden drop impact and its role in disease spread.

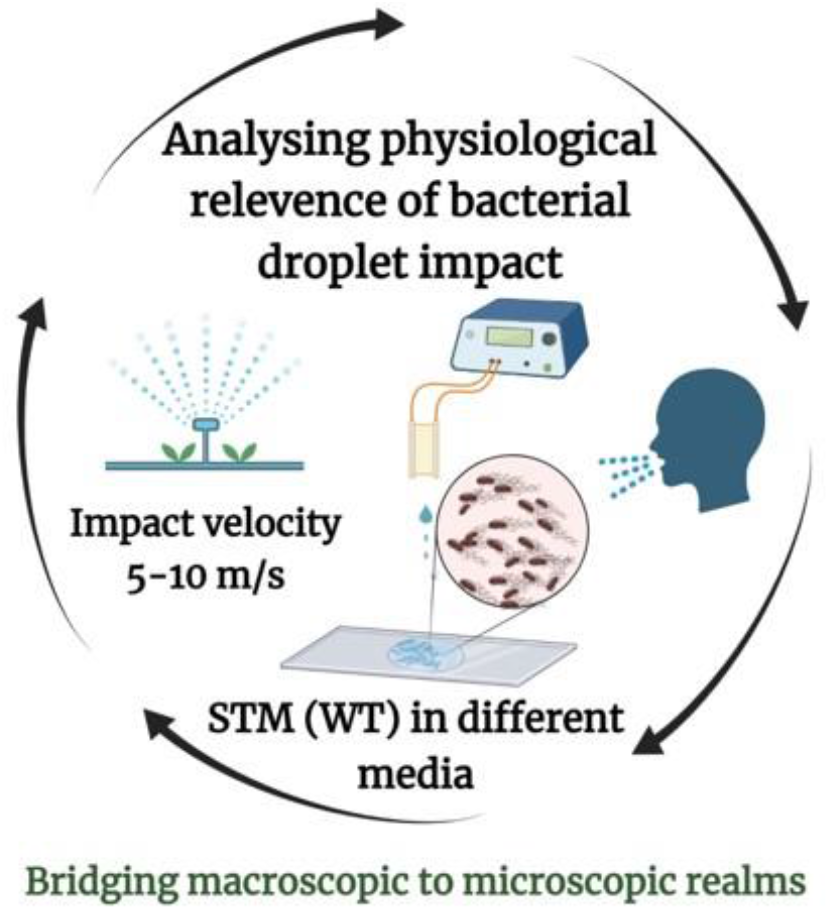

---

The liquid droplet impact on surfaces plays an important role in various natural processes. In general terms, human respiratory activities expel droplets during talking, coughing, sneezing practices and droplet impact is crucial in various engineering applications. The impact dynamics profoundly depend on the physicochemical characteristics of impact substrate and droplet properties. The characterization of droplet impact is carried out with the aid of several dimensionless numbers: the Reynolds number (Re=ρDV/μ), which is a dimensionless ratio of inertial to viscous force Weber number (We = ρDV^2^γ), which is the ratio of inertia to surface tension forces (γ), Ohnesorge number 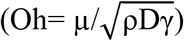, and Capillary number (Ca=μV/γ) where μ is the viscosity, ρ is the density, D is the diameter, γ is the surface tension forces and V is the impact velocity. However, the experimental investigation realm on droplet impact is confined only to the physical aspect of droplet dynamics for a lower range of Weber numbers. The droplet impact mechanism for water drops impacted at exceptionally low velocities was primarily reported by Worthington^1^. The literature reports substantial work on droplet spreading, receding, splashing, and bouncing^2–5^. Researchers have explicitly mentioned that the bacterial laden droplet leads to potential colonization and the ability to transmit fomites to humans irrespective of the droplet size and the surface it falls. The respiratory droplets emitted during exhalation, coughing and sneezing span in-between a wide range of droplet size and velocity scales (Supplementary Figure 1). The microdroplets have a modest Stokes number that can be impacted by ambient airflow convection, and the droplets transport much farther than 1.83m ^6^. However, such microdroplets impacting a surface may pose potential risks to people when they touch such surfaces by chance. In this context, the present study deals with the microscopic size bacteria-laden drops that impact at a certain weber number onto a surface and to understand the relentless effect of the deposited bacterial patterns for their virulence and bacterial infectivity at different velocities.

The bacterial droplet impact is an unexplored area of research but has a profound relevance that requires unequivocal demonstration to comprehend the impact dynamics and effects on bacteria subjected to a broader range of impact velocity. Environmental forces have an obvious impact on the phenotypes of bacteria or any other microscopic organisms. As a result, stress-induced mechanical forces influence bacterial physiology, which is poorly understood. However, each bacteria within the medium can experience contact force when it gets acquainted with any surface. These contact forces can either be compressive or tensile, depending on whether the bacterial cell wall pushes itself onto or pulled away from the surface. Further, they experience compressive forces when exposed to the open atmosphere or deep-sea water environments when inundated by some host cells. *Salmonella* is a common bacterial type that severely impacts the feed supply chain and various food production industries^7^. It has been reported that thousands of people die worldwide from *Salmonella* systemic infections like non-typhoidal salmonellosis (NTS), brain culminating in meningitis, and associated neurological abnormalities ^8–12^. Moreover, *Salmonella* continues to be the second biggest cause of foodborne disease in the United States and the leading cause of both hospitalization and deaths ^7^. *Salmonella* infection spreads to healthy individuals through contaminated food and water ^13^. However, airway transmission of different *Salmonella enterica* serovars such as Enteritidis, Typhimurium, and Agona among poultry animals has also been reported ^14,15^.

Aerosolized *Salmonella* can colonize in the ceca, trachea, liver, and spleen of poultry broilers^16^. Despite their poor *in vitro* survival due to evaporation stress *Salmonella* Typhimurium (STM) retrieved from inanimate surfaces showed hyper-virulence in RAW264.7 cells ^17^. These aspects add to the importance of studying the effect of *Salmonella*-laden drops for various impact velocities and base fluid mediums (nutrient-rich or neutral) to understand the transmission risk.

A recent study indicates that the evaporation of droplets can result in respiratory aerosols that can drive the transmission of SARS-CoV-2^18^. In this regard, droplet impact studies have immense importance from the perspective of virulence and the infectivity of the bacteria-laden droplets for varying impact velocities embedded onto a surface. Subsequently, the focussed research in the regime of 5-10 m/s impact velocities is also crucial, which mimics the velocity range of the droplets expelled from the mouth and nose during the sneezing or coughing process. Henceforth, the physiological relevance of bacterial droplet impact is unequivocally studied in this investigation for the impact velocities of 5-10 m/s. Our goal is to understand how the visible effects are related to the microscopic behavior of bacteria-laden drop impact. Figure 1 (a) represents the schematic diagram for the instrumentation for measuring impact velocity, spreading diameters, and the impact droplet deposition patterns captured using side-view imaging, fluorescence microscopy, and reflection interference microscopy. Further, the bacterial viability for the impacted velocity ranging from 5-10 m/s, the infectivity of the impacted droplets, and the effect of the RNA isolation process are studied as depicted in Figure 1(b).

**Fig.1.**
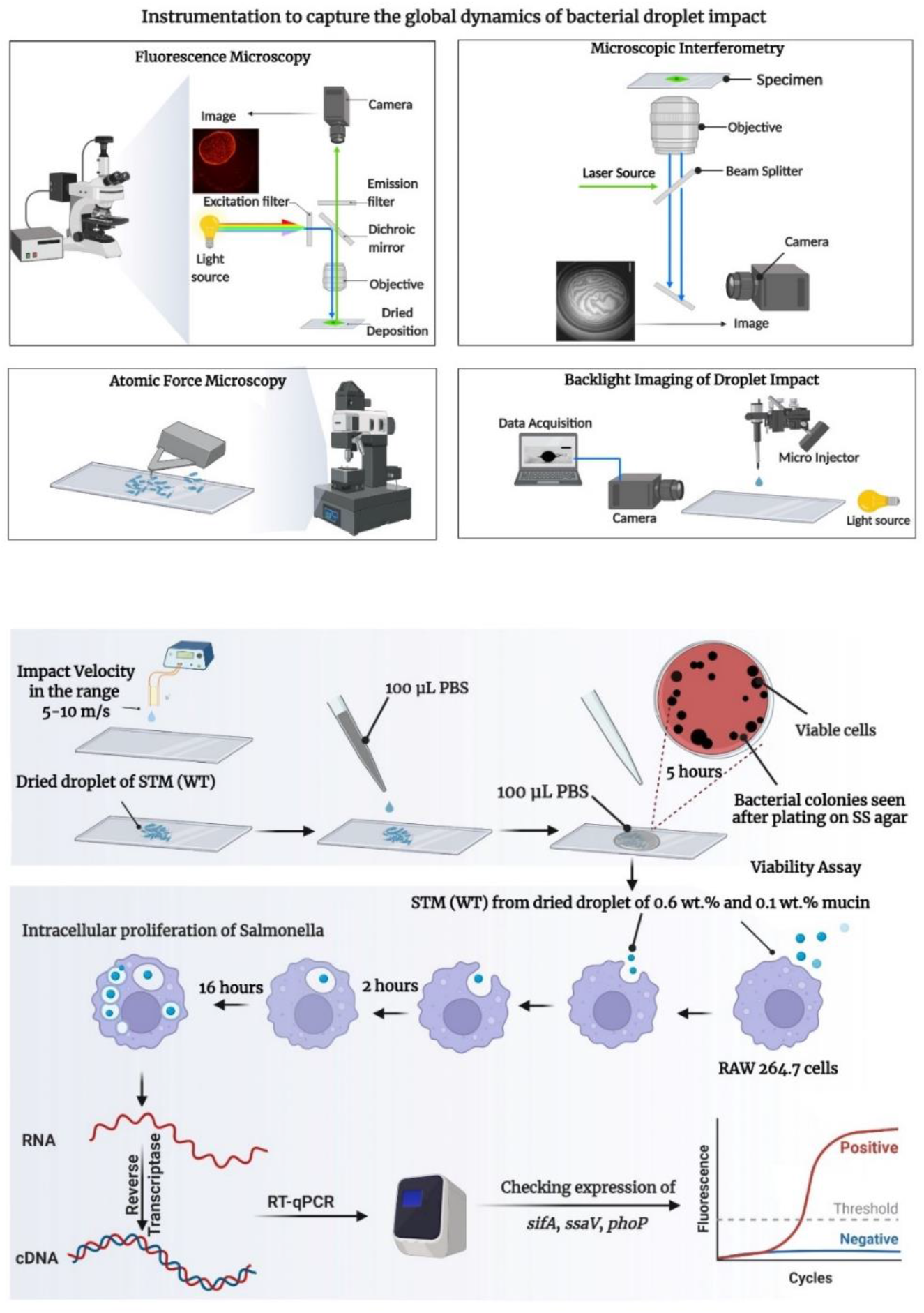
Schematic layout of the experimental study. **(a)** The schematic representation to capture the global dynamics of droplet impact involving fluorescence microscopy, high-speed microscopic interferometry, atomic force microscopy and backlight imaging of bacterial droplet impact. **(b)** Experimental layout depicting the diagnosis of physiological relevance of bacterial drop impact involving in-vitro studies, viability assay and RNA isolation. Diagram created with BioRender (www.biorender.com).

## Global Impact Dynamics of bacteria-laden droplets signifies the dominance of inertial effects

The droplet spread over the glass substrate primarily depends on the inertial effects, based on the Weber number corresponding to the impact velocities of 5,7 and 10 m/s. The side-view imaging of the droplet impact on the glass surface is presented in Figure 2(a) at various time instants. Droplet impact sequence from the point of contact on the surface to the extent of spreading and to retraction towards the formation of a sessile drop (i.e., for a higher velocity of 10 m/s) is evident from the time instants. Figure 2(b) represents the spreading dynamics of the droplet at a higher impact velocity and the variation of thin-film liquid layers captured using reflection interference microscopy. The droplet spreading is entrenched with various parameters such as viscous dissipation, surface tension effects, surface roughness, and contact line dynamics. These factors get overshadowed when the inertia forces overcome the effects caused due to viscosity.

**Fig.2.**
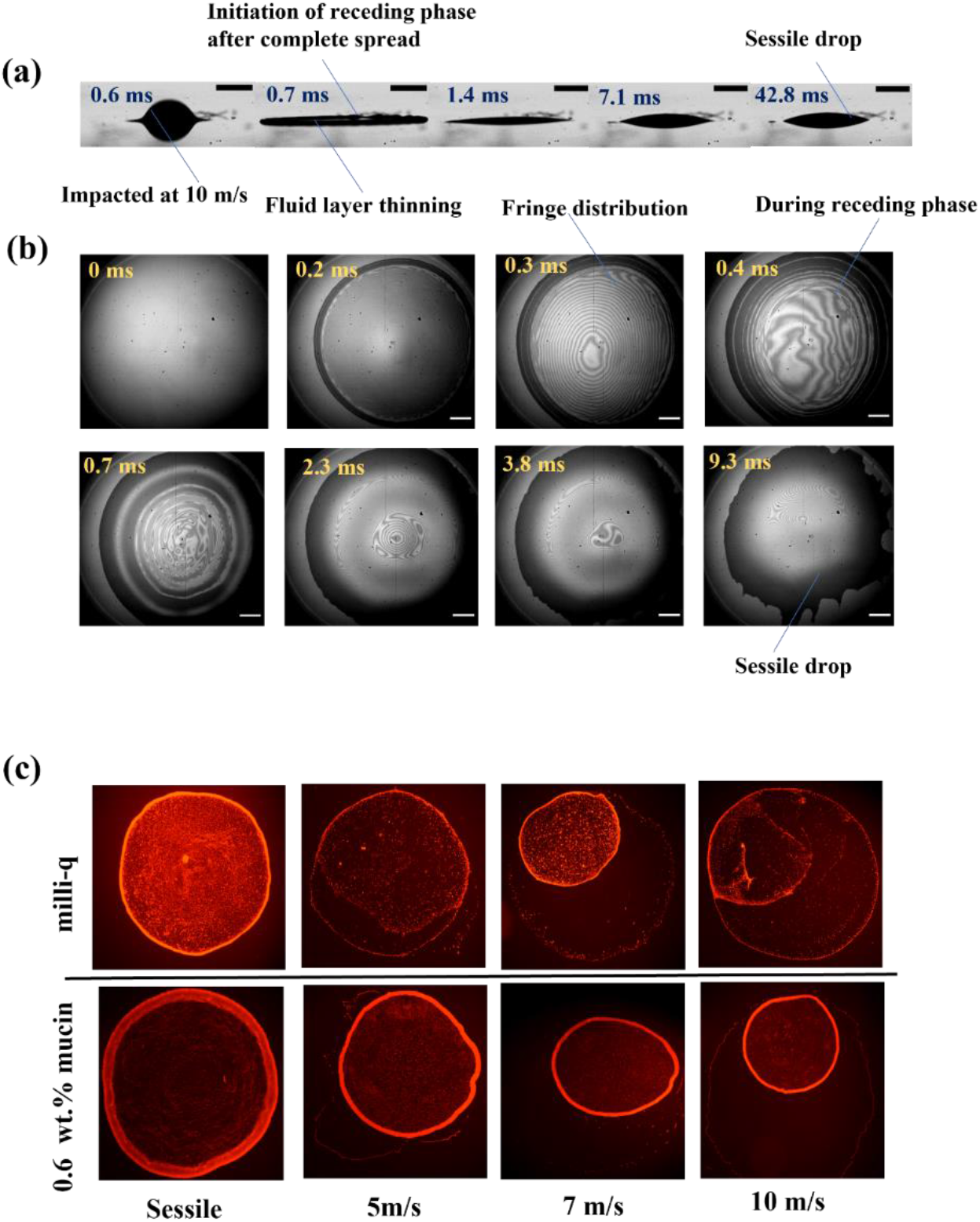
Global dynamics of bacterial droplet deposition. **(a)** Profile view Images captured at 10000 fps for Salmonella laden droplet impacted on glass surface with 10 m/s on the glass substrate. Scale bar is 400 μm. **(b)** High-speed interference images captured at 10000 fps for Salmonella laden droplet impacted on glass surface with 10 m/s on the glass substrate. Scale bar is 200 μm. **(c)** Fluorescence intensity captured for dried droplet impacted at different Weber number

In this context, the present study demonstrates the influence of inertia forces rather than viscous effects during the bacterial droplet impact and spreading. The bacterial droplet spreads after the first point of contact on the surface and reaches a maximum spreading diameter (*d_max_*) until the initial kinetic energy is absorbed entirely. The spreading time also decreases with increased impact velocity, as the higher kinetic energy is embedded in the droplet. Further, when the drop attains *d_max_* the bacteria-laden drop retracts as the impact weber number is significant, for which the maximum spreading diameter *d_max_* is always greater than the equilibrium spreading diameter. Similarly, a higher receding velocity is observed for the higher impact velocity, as shown in Table 1. The interference patterns observed during the spread of the liquid layer signify the presence of a relatively thin liquid layer during the initial time instants for the maximum spread in the droplet. As time progresses, retraction of the liquid layer is evident from the recorded images. Hence, the propagation of capillary waves of the receding liquid layer is seen in the form of dark circular bands in the images from *t* = 0.3 to 0.7 ms (Fig.2b).

**Table 1.**
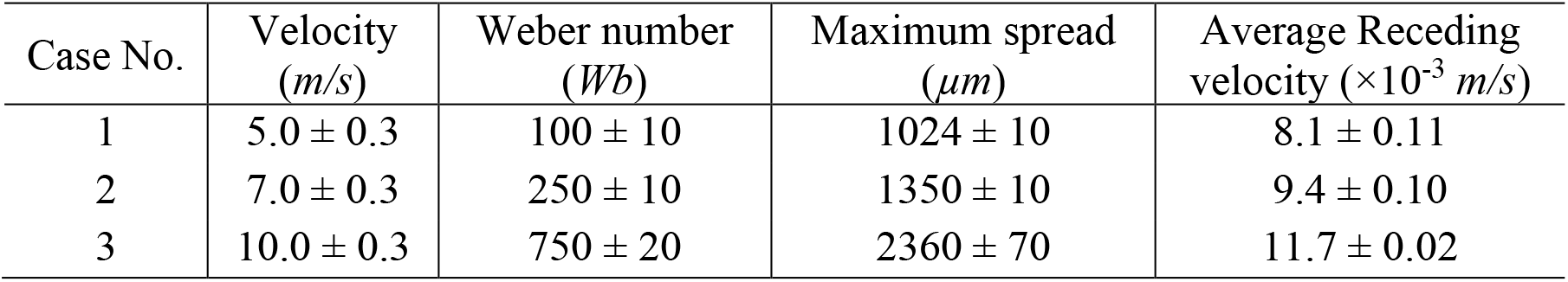
Operating conditions on which the experiments are carried out indicating the impact velocities along with maximum spread and diameter of drop investigated.

| Case No. | Velocity<br>(*m/s*) | Weber number<br>(*Wb*) | Maximum spread<br>($\mu m$) | Average Receding<br>velocity ($\times 10^{-3}$ *m/s*) |
| --- | --- | --- | --- | --- |
| 1 | $5.0 \pm 0.3$ | $100 \pm 10$ | $1024 \pm 10$ | $8.1 \pm 0.11$ |
| 2 | $7.0 \pm 0.3$ | $250 \pm 10$ | $1350 \pm 10$ | $9.4 \pm 0.10$ |
| 3 | $10.0 \pm 0.3$ | $750 \pm 20$ | $2360 \pm 70$ | $11.7 \pm 0.02$ |

**Table 2.** Strains used in this study.

| Strains/ plasmids | Characteristics | Source/ references |
| --- | --- | --- |
| *Salmonella enterica* serovar Typhimurium ATCC strain 14028S | Wild-type (WT) | Gifted by Prof. M. Hensel |
| *S. Typhimurium* Δ*phoP* | Chl<sup>R</sup> | Laboratory stock |

**Table 3.**
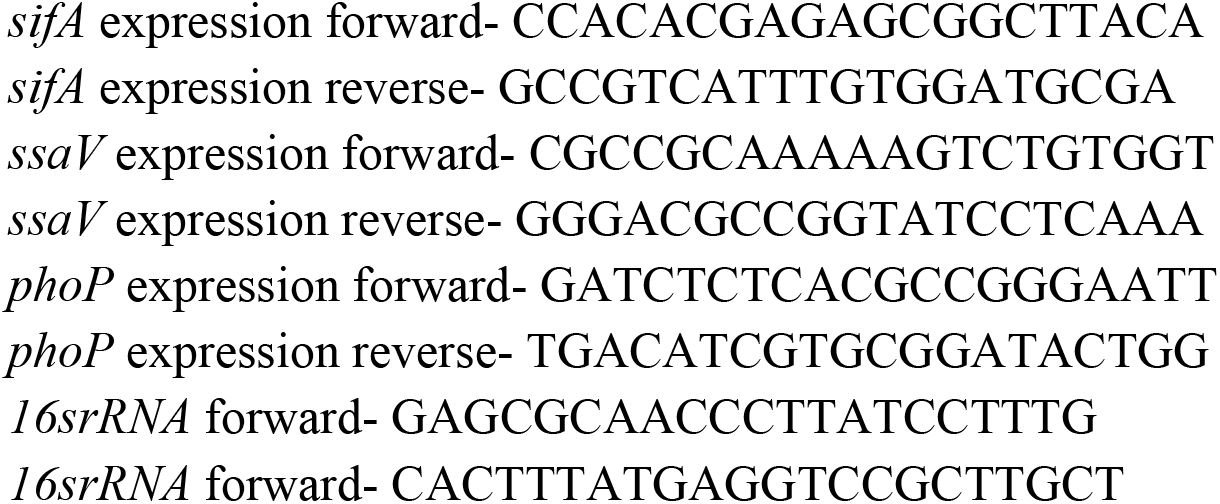
Oligos used in this study.

The increase in multiple waves can be seen in the receding process of the liquid layer and disintegration of the waves is seen once it reaches the form of a sessile drop. The fringe patterns during the spread process are due to the thin-film liquid layer at lower droplet contact angles. The distribution of fringe pattern is relayed on the increase in contact angle from the receding point to the formation of sessile drop (Fig.2b). The global dynamics of the droplet spread almost remain similar for the various inertial forces, where the viscous forces do not have a significant effect (Supplementary Figure 2). The side-view and interference microscopy imaging have been performed simultaneously with high-speed imaging systems as the impact, spreading, and retraction occurs in the rapid alteration regimes. The increase in the impact velocity of the bacteria-laden droplet leads to a more extensive spread of the liquid layer, as presented in Table 1, due to an increase in inertial force. Understanding the droplet impact dynamics assists in determining the stress experienced by the bacteria during the impact process, and their possible effect on the bacterial morphology is discussed further.

The drying effects of bacteria-laden sessile drops after impact on the surface are inadequately addressed in the literature. The evaporation of impacted droplets leaves a different dried pattern on the glass substrate compared to sessile deposition. It is observed that the deposited pattern can range from a simple ring-like structure which is referred to as a coffee ring, or multiple coffee rings with uniform deposition, depending on the shear force and flow characteristics. Correspondingly, the bacteria-laden fluids revealed a variety of patterns on the glass substrate after the impacted drop evaporated, as presented in Fig.2(c). A thick outer ring of particles was seen in the dried biological STM-mucin samples. For a sessile drop, the fluid flows radially outwards to replace the evaporated fluid at the base edge of the drop, maintaining a constant radius and thus leading to a commonly seen coffee ring effect ^19,20^. Moreover, capillary flow creates a thick outer ring at its maximum spread when the droplet is impacted at a certain velocity for which the viscosity of the base fluid medium is accountable for carrying the particle towards the contact line ^21^. Further, when the droplet recedes after reaching maximum spread, it experiences a constant contact angle regime, and the bacterial deposition was found to be uniform inside the ring. The variation in deposition patterns primarily indicates differences in bacterial morphology with increase in impact velocities which is systematically investigated in the following sections.

## Varying stress conditions immensely altered the bacterial morphology

From the discussions in the earlier section, inertial force is found to have a significant role in altering bacterial morphology. The topographic images of the bacterial cell wall at three impact weber numbers (100,250 and 750) corresponding to 5,7 and 10 m/s are acquired using Atomic Force Microscopy (AFM) in nutrient neutral (milli-Q) and nutrient-rich medium (0.6 wt.% mucin). The coupling of mechanical forces with microscopic characterization is essential for comprehending physiological changes. The qualitative analysis of the droplet impact with rapidly changing conditions has a significant effect on the surface morphology of the bacteria. Like other Gram-negative pathogens, *Salmonella* has a rigid cell wall made of peptidoglycan, a repeating unit of N-acetyl muramic acid and N-acetyl glucosamine connected by pentapeptide linkage ^22,23^. In response to environmental stimuli, *Salmonella* modifies its peptidoglycan layer ^24^. When subjected to a certain velocity, the shape regulation of the *Salmonella* cell wall is profoundly affected by the external forces that affect the rate of cell wall synthesis^25^. Apart from the impact stress, the bacterial cell wall is subjected to the evaporative stress, which does not allow the bacteria to snap back to its initial cell wall shape (Fig.3a) as vividly observed for the Wb=250 (5m/s) and 750 (7m/s) for milli-Q and nutrient-rich medium^26^ (Fig. 3b,3c and 3d).

**Fig.3.**
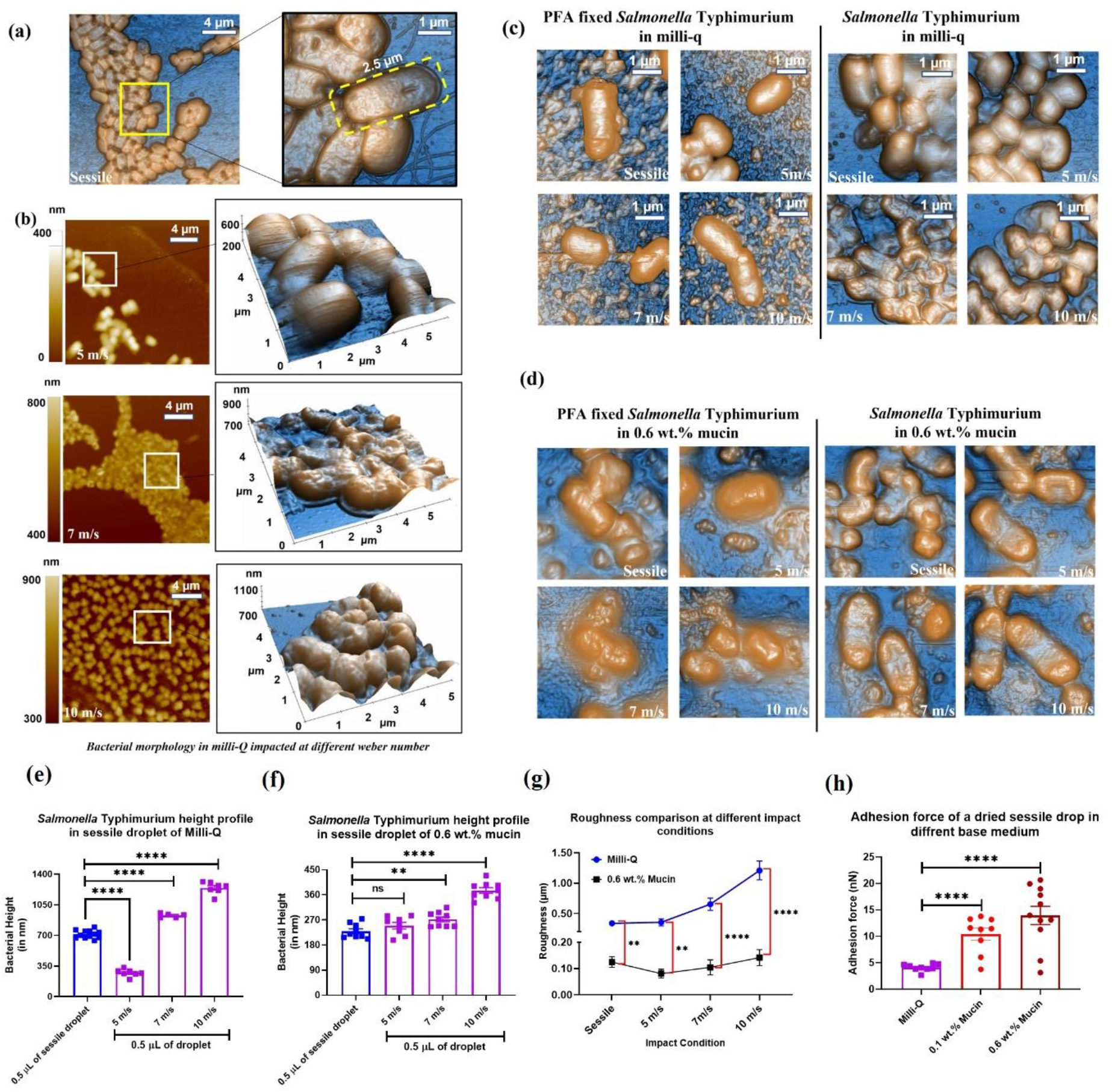
Atomic Force Microscopy identifies larger stress induction and greater surface damage at higher impact velocity. **(a)** AFM images for *Salmonella* in milli-Q for a sessile drop **(b)** AFM images of bacterial laden droplet with milli-Q as base medium at three different impact velocities in 5μm square area **(c)** AFM images of bacterial laden droplet with milli-Q and **(d)** 0.6 wt.% mucin as base medium compared with PFA fixed bacteria at three different impact velocities along with their height profiles in 5μm square area **(e)** Adhesion energy in nutrient neutral (milli-Q) and nutrient-rich (0.6 wt.% Mucin) base fluid medium experienced on a dried STM (WT) sessile drop **(f)** Roughness measurements presented as RMS roughness at different impact conditions for milli-Q and 0.6 wt.% mucin. ***(P)* *< 0.05, *(P)* **< 0.005, *(P)* ***< 0.0005, *(P)* ****< 0.0001, ns= non-significant, (Student’s t test-unpaired)**

The maximum bacterial height of *Salmonella* when subjected to desiccation stress and the mechanical stress induced due to the impact velocity increased from 712.6 (sessile) to 1244.3 nm (10 m/s) in milli-Q medium, as shown in Fig.3(e). A similar trend is seen when milli-Q is replaced with a nutrient-rich, 0.6 wt.% mucin solution (Fig.3f), indicating flexible behavior of the cell walls as seen in Fig.3(c) and (d). AFM data also supports the view that the cell walls of *Salmonella* Typhimurium deform flexibly in the milli-Q and mucin medium, as shown in Fig.3(c), due to nutrient deficiency and evaporative stress. Moreover, shear stress is induced in the cell walls of the bacteria at higher impact velocities (Supplementary Figure 3). The height profile of a randomly selected bacterial cell (Supplementary Figure 4) indicates the presence of peaks and valleys over the cell wall. The deformation in the cell wall creates peaks and valleys can be quantified using the roughness parameter.

The root mean square roughness (*R_q_*) is calculated for *Salmonella* Typhimurium milli-Q and 0.6 wt.% mucin is calculated based on the following Eq. (1) for sessile and impact conditions (5,7 and 10 m/s).

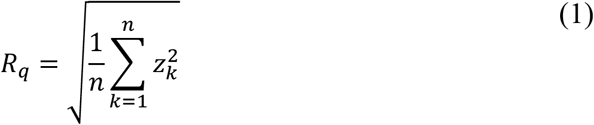

where, *R_q_* is the root mean square roughness at a height ‘*z*’ for the pixel ‘*k*’ in the image. The *R_q_* indicates the distance between peaks and valleys or the average height variations from the mean line. Physically, *R_q_* demonstrates the extend of surface damage which is significant in milli-Q than in 0.6 wt. % mucin for Weber number greater than 250 (5 m/s) as shown in Fig. 3(g). The stress experienced on the deposition surface, calculated from the force exerted by the AFM probe, shows higher surface stress (Supplementary Figure 3) for velocity greater than 7 m/s with increased surface roughness. The nano-intender used for Atomic Force Microscopy is used to acquire the adhesion energy for nutrient-rich and nutrient-neutral samples by quantifying the forces between the tip and the sample.

The force-displacement spectroscopy for all the samples indicates that the force experienced during intending on the dried bacterial sample during approach and retraction are significantly different, depicting greater adhesion between the tip and the cell walls, especially for velocity greater than 7m/s (Fig. 3h). Furthermore, higher adhesion energy indicates enhanced interaction of cells with the adjacent cells and the substrate on which the impact occurs. The enhanced cell-cell interaction leads to local stiffening of the dried deposit resulting in variations in the adhesion energy and resisting the deformation caused due to the receding velocity at different impact conditions. This enhanced interaction between the cells and substrate cushions the bacteria, keeping it viable even in adverse conditions^27^. The increased surface roughness has a pertinent role in microbial adhesion^28,29^. The undulations on the surface of bacterial deposits caused due to surface roughness enhanced their adhesion on glass substrate due to increased surface area and fissures at higher impact velocities. Moreover, cell-cell interaction or cushioning of bacteria happens when the adhesion energy is large, especially in the case of nutrient-rich 0.6 wt.% mucin compared to a nutrient-deficient milli-Q medium (Fig.3g).

The cushioning effect on the cell membranes is quantitatively assessed from the roughness kurtosis. The roughness kurtosis (*R_ku_*), acquired from AFM, provides the sharpness of spikes on the surface, which physically denotes that a grooved or pitted surface could shelter the bacteria by providing a cushioning effect. The numerical value of *R_ku_* greater than three indicates a presence of a spiky surface, and *R_ku_* less than three denotes a bumpy surface^30^ given by Eq.(2).

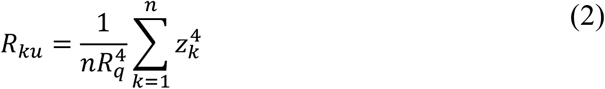

where *R_q_* is the root mean square roughness at a height ‘*z*’ for the pixel ‘*k*’ in the image. The average roughness kurtosis is ≈5.1 in the case of bacteria-laden 0.6 wt.% mucin, ≈4.2 in 0.1 wt.% mucin and ≈1.1 in milli-Q sessile drop. The spiky and bumpy surfaces are evident from the AFM images, as shown in Figure 3(a),3(c) and 3(d). This indicates that mucin (0.1 and 0.6 wt%) provides a better cushioning effect to the bacteria trapped between the spiky surfaces than milli-Q and may influence the *in vitro* viability of the bacteria. However, the viability of the bacteria is subjected to nutrient availability in the medium, which requires further investigation, as discussed in the subsequent sections.

## The velocity reduced the *in vitro* viability of *Salmonella* Typhimurium in nutrient-rich desiccated droplets

Earlier, we reported that irrespective of the presence of nutrients, the *in vitro* viability of wild-type *Salmonella* inside the sessile droplets is severely compromised post-evaporation. Moreover, the bacteria recovered from dried nutrient-rich sessile droplets hyper-proliferates inside the RAW264.7 macrophages ^17^.

However, the *in vitro* viability of bacteria retrieved from the impacted dried droplets on solid surfaces with specific velocities has not been tested before. In this study, we have used neutral (milli-Q), low-nutrient (glucose 5 wt%, and 10% glycerol), and nutrient-rich (LB broth, mucin 0.1 wt.%, and 0.6 wt.%) base solutions for impacting the bacterial droplet on the glass surface with specific velocities (5, 7, and 10 m/s) as mentioned earlier. 0.5 μL of the planktonic culture of wild-type *Salmonella* in different base solutions gave rise to 10^5^ to 10^6^ CFU of viable bacteria (Fig. 4a– 4f; red bar). In the dried sessile droplet of 0.5 μL volume, the viability of STM (WT) reduced drastically to 10^2^ to 10^4^ CFU (Fig. 4a– 4f; blue bar). The loss of moisture due to evaporation restricted the availability of nutrients to the bacteria in the dried droplet and decreased the bacterial burden compared to the liquid-phase culture. In milli-Q (Fig. 4a) and 5% glucose solutions (Fig. 4b), the impact of wild-type *Salmonella* droplet on glass surface did not create any significant difference in its viability (~10^2^ CFU) compared to the sessile droplet (~10^2^ CFU). 0.5 μL of the planktonic culture of wild-type *Salmonella* in different base solutions gave rise to 10^5^ to 10^6^ CFU of viable bacteria (Fig. 4a– 4f; red bar).

**Fig.4.**
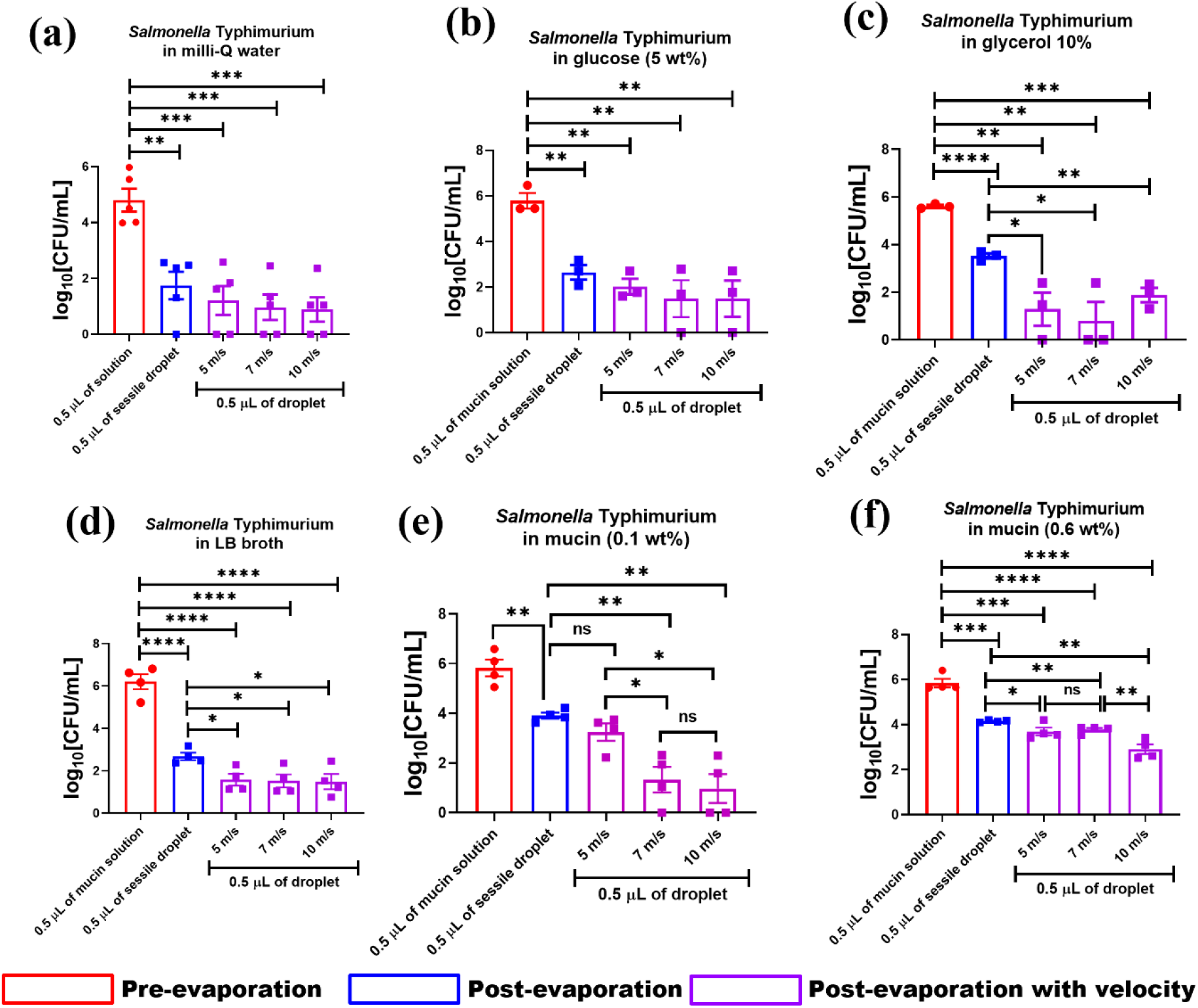
Assessment of *in vitro* viability of *Salmonella* retrieved from desiccated droplets of different base solutions impacted on a solid surface with or without velocities. The viability of *Salmonella* Typhimurium (in log10 scale) recovered from dried sessile droplets of different base solutions impacted on solid glass surface with or without velocities (5, 7, and 10 m/s). As base solutions **(a)** milli-Q, **(b)** glucose 5 wt%, **(d)** 10% glycerol, **(e)** LB broth, and **(f)** mucin 0.1 wt%, were used **(N≥3)**. Data are represented as mean ± SEM. ***(P)* *< 0.05, *(P)* **< 0.005, *(P)* ***< 0.0005, *(P)* ****< 0.0001, ns= non-significant, (Student’s t test-unpaired)**

In the dried sessile droplet of 0.5 μL volume, the viability of STM (WT) reduced drastically to 10^2^ to 10^4^ CFU (Fig. 4a– 4f; blue bar). The loss of moisture due to evaporation restricted the availability of nutrients to the bacteria in the dried droplet and decreased the bacterial burden compared to the liquid-phase culture. In milli-Q (Fig. 4a) and 5% glucose solutions (Fig. 4b), the impact of wild-type *Salmonella* droplet on glass surface did not create any significant difference in its viability (~10^2^ CFU) compared to the sessile droplet (~10^2^ CFU). In milli-Q (Fig. 4a) and 5% glucose solutions (Fig. 4b), the impaction of wild-type *Salmonella* with glass surface did not create any significant difference in its viability (~10^2^ CFU) compared to the sessile droplet (~10^2^ CFU). *Salmonella* can survive in glycerol by upregulating the genes associated with lipopolyssacharide biosynthesis and gluconate metabolism^31^. When the bacteria were allowed to impact the glass slide with glycerol (10%) as the medium (Fig. 4c), we observed a significant decrease in the *in vitro* viability of *Salmonella* (~10^2^ CFU) compared to the sessile droplet (~10^4^ CFU) under all three velocities (Fig. 4c).

The reduction in the bacterial viability under velocity can be attributed to mechanical stress-induced due to impact velocity (Supplementary Figure 3). Similar observations were obtained when bacteria-laden nutrient-rich-based solutions such as LB broth (Fig. 4d), 0.1 wt%, and 0.6 wt% mucins (Fig. 4e and 4f) were subjected to impact velocity. The viability of STM (WT) in the sessile droplet (~10^3^ CFU) of LB broth was significantly reduced to ~10^2^ CFU when exposed to impact velocity (Fig. 4d). A more prominent velocity-induced reduction of *in vitro* viability was observed in 0.1 wt% mucin solution (Fig. 4e). Mucins are heavily glycosylated O-linked glycoproteins, synthesized and secreted by the epithelial cells ^32^. The availability of nutrients in mucin improved the survival of the bacteria in the sessile droplet (~10^4^ CFU) even when subjected to an impact velocity of 5 m/s (~10^4^ CFU) velocity (Fig. 4e).

With an increase in the impact velocity from 5 to 7 m/s, the viability of wild-type *Salmonella* decreased from ~10^4^ CFU to ~10^2^ CFU and remained constant at 10 m/s (~10^2^ CFU) (Fig. 4e). Further, when we used 0.6 wt% of mucin, we observed that the higher nutrient content overshadowed the effect of impact velocity on the *in vitro* survival of *Salmonella* (Fig. 4f). Unlike 0.1 wt% mucin concentration, the reduction in the viability of STM (WT) was observed between 7 (~10^4^ CFU) and 10 m/s (~10^3^ CFU), as depicted in Fig.4(f). Collectively the present data suggest that in addition to desiccation and nutrient limitation stress, the mechanical stress induced by the impact velocity also reduces the bacterial burden in dried droplets post-evaporation.

## Wild-type *Salmonella* from the desiccated droplet of mucin subjected to higher impact velocity indicated hyperproliferation in RAW264.7 cells

We further investigated the effect of velocity-induced mechanical stress in the intracellular virulence of *Salmonella* retrieved from the dried droplet of mucin.

After crossing the intestinal mucosal barrier, Salmonella interacts with phagocytic immune cells such as macrophages, dendritic cells, neutrophils, etc.^13^. Professional antigen-presenting cells like macrophages phagocytose invading pathogens limit their spread throughout the host body^33,34^. Our study revealed that Salmonella recovered from the dried sessile and velocity impacted mucin (0.1 wt%) droplets are prone to phagocytosis by RAW264.7 cells compared to the planktonic culture (Fig 5a), suggesting a better clearance of the bacteria and subsequent restriction of the infection by macrophages. Surprisingly, we found these phagocytosed bacteria hyper-proliferate inside the macrophages (Fig 5b), indicating that the bacteria droplet subjected to a specific impact velocity efficiently maintains their virulence while infecting the host cells. When the Salmonella recovered from the post-evaporated dried droplets of high mucin concentration (0.6 wt%) and used to infect the RAW264.7 cells, we noticed similar phenotypes (Supplementary Figure-5a and 5b). Inside the host cells, Salmonella stays inside a membrane-bound acidic compartment called Salmonella containing vacuole (SCV)^35^. The acidic pH of SCV is sensed by the PhopQ two-component system (TCS) of Salmonella, which uses Salmonella pathogenicity island-2 (SPI-2) encoded virulent factors such as sifA, ssaV, etc. for its successful replication inside the host cell^36–38^.

**Fig.5.**
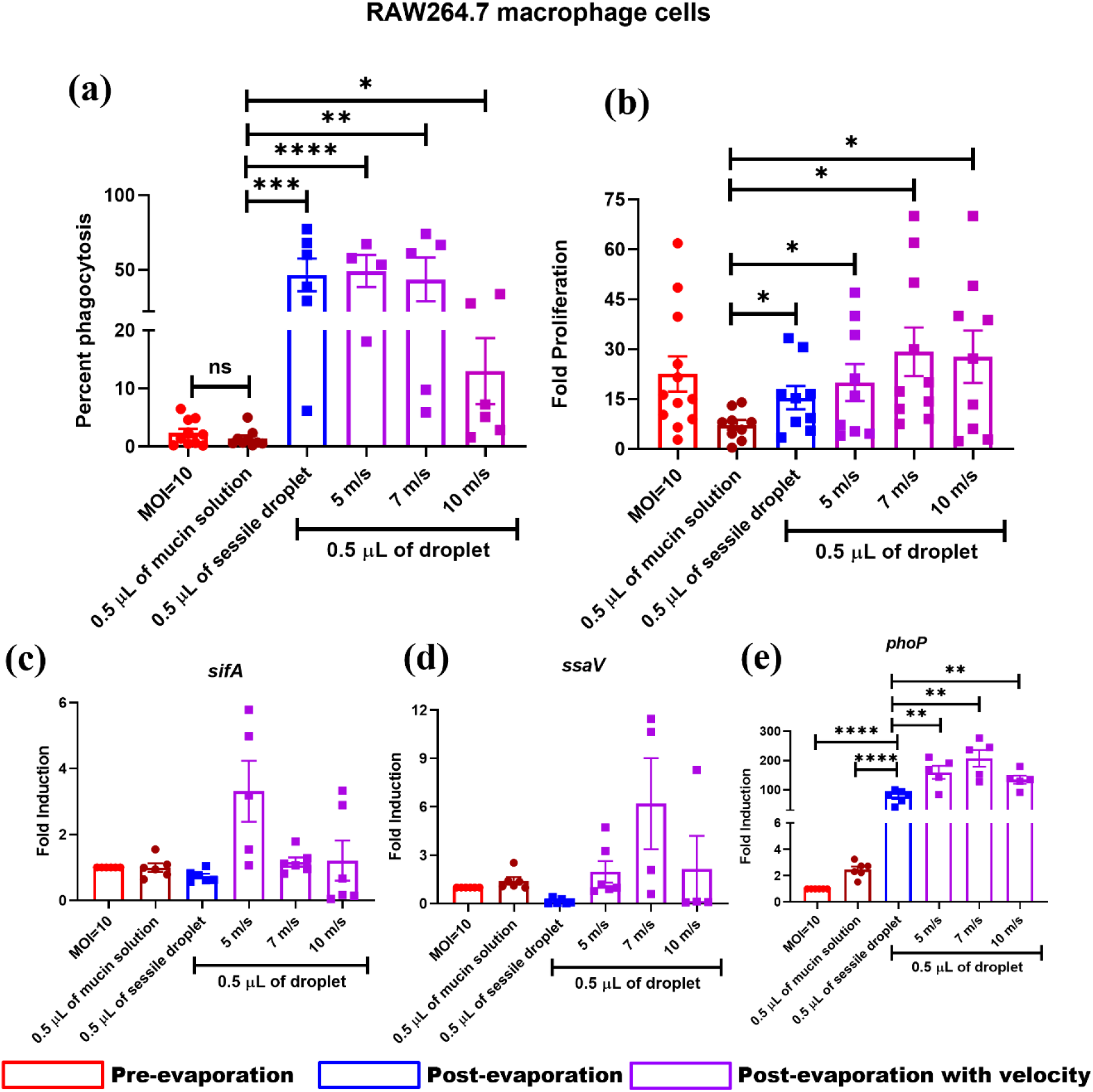
The virulence of *Salmonella* Typhimurium retrieved from desiccated mucin droplet (0.1 wt.%) impacted on a solid surface at different velocities and for the sessile droplet. *Salmonella* Typhimurium recovered from desiccated mucin (0.1 wt%) droplet impacted solid glass surface with or without specific velocities (5, 7, and 10 m/s) were used to infect RAW264.7 cells to determine the **(a)** percent phagocytosis and **(b)** intracellular proliferation (n=3, N=2). Determining the transcript-level expression of **(c)** *sifA*, **(d)** *ssaV* and **(e)***phoP* from intracellular *Salmonella* by RT-qPCR (n=3, N=2). Data are represented as mean ± SEM. ***(P)* *< 0.05, *(P)* **< 0.005, *(P)* ***< 0.0005, *(P)* ****< 0.0001, ns= non-significant, (Student’s t test-unpaired)**

The present study, hypothesized that the velocity-induced mechanical stress upregulated the expression of PhopQ TCS and SPI-2 genes in Salmonella, which help in the successful proliferation of the bacteria inside the macrophages. With an increase in the impact velocity (5 to 10 m/s) of Salmonella retrieved from 0.1 (Fig. 5c–5e) and 0.6 wt% (Supplementary Figure 5c-5e) mucin droplets, the transcript-level expression of sifA (Fig. 5c), ssaV (Fig. 5d), and phoP (Fig. 5e) increased in intracellular bacteria, which explains the reason behind their enhanced proliferation inside the macrophages. Surprisingly, we found these phagocytosed bacteria hyper-proliferate inside the macrophages (Fig 5b), indicating that the bacteria droplet subjected to a specific impact velocity efficiently maintains their virulence while infecting the host cells. When the Salmonella recovered from the post-evaporated dried droplets of high mucin concentration (0.6 wt%) and used it to infect the RAW264.7 cells, we noticed similar phenotypes (Supplementary Figure-5a and 5b). Inside the host cells, Salmonella stays inside a membrane-bound acidic compartment called Salmonella containing vacuole (SCV)^35^. The acidic pH of SCV is sensed by the PhopQ two-component system (TCS) of Salmonella, which uses Salmonella pathogenicity island-2 (SPI-2) encoded virulent factors such as sifA, ssaV, etc. for its successful replication inside the host cell^36–38^. We hypothesized that the velocity-induced mechanical stress upregulated the expression of PhopQ TCS and SPI-2 genes in Salmonella, which help in the successful proliferation of the bacteria inside the macrophages. With an increase in the impact velocity (5 to 10 m/s) of Salmonella retrieved from 0.1 (Fig. 5c–5e) and 0.6 wt% (Supplementary Fig. 5c-5e) mucin droplets, the transcript-level expression of sifA (Fig. 5c), ssaV (Fig. 5d), and phoP (Fig. 5e) increased in intracellular bacteria, which explains the reason behind their enhanced proliferation inside the macrophages.

## Survival of wild-type *Salmonella* in the desiccated mucin droplets depends upon *phoP*

As wild-type *Salmonella* recovered from the dried droplet of mucin (for sessile and impact velocities 5,7 and 10 m/s) indicated a consistent upregulation in the expression of *phoP* while infecting macrophages, we hypothesized that the survival of *Salmonella* Typhimurium in the nutrient restricted environment of fomites depends upon *phoP*. The assessment has been carried out for *in vitro* viability of STM (WT) (Fig. 6a) and *ΔphoP* (Fig. 6b) in sessile droplets of high concentration mucin (0.6 wt%) to the impact Weber numbers of 100, 250 and 750. As discussed earlier, the viability of STM (WT) (~10^6^ CFU to ~10^4^ CFU) and *ΔphoP* (~10^5^ CFU to ~10^3^ CFU) was compromised in the sessile droplet of mucin after evaporation (Fig. 6a and 6b). Impaction of mucin solutions on the glass surface with 5, 7, and 10 m/s velocities further reduced the viability of STM (WT) (~10^2^ CFU) due to the generation of mechanical stress (Fig. 6a). On the contrary, STM *ΔphoP* struggled to survive in the desiccated mucin droplets for the impact Weber numbers 100-750 (Fig. 6b).

**Fig.6.**
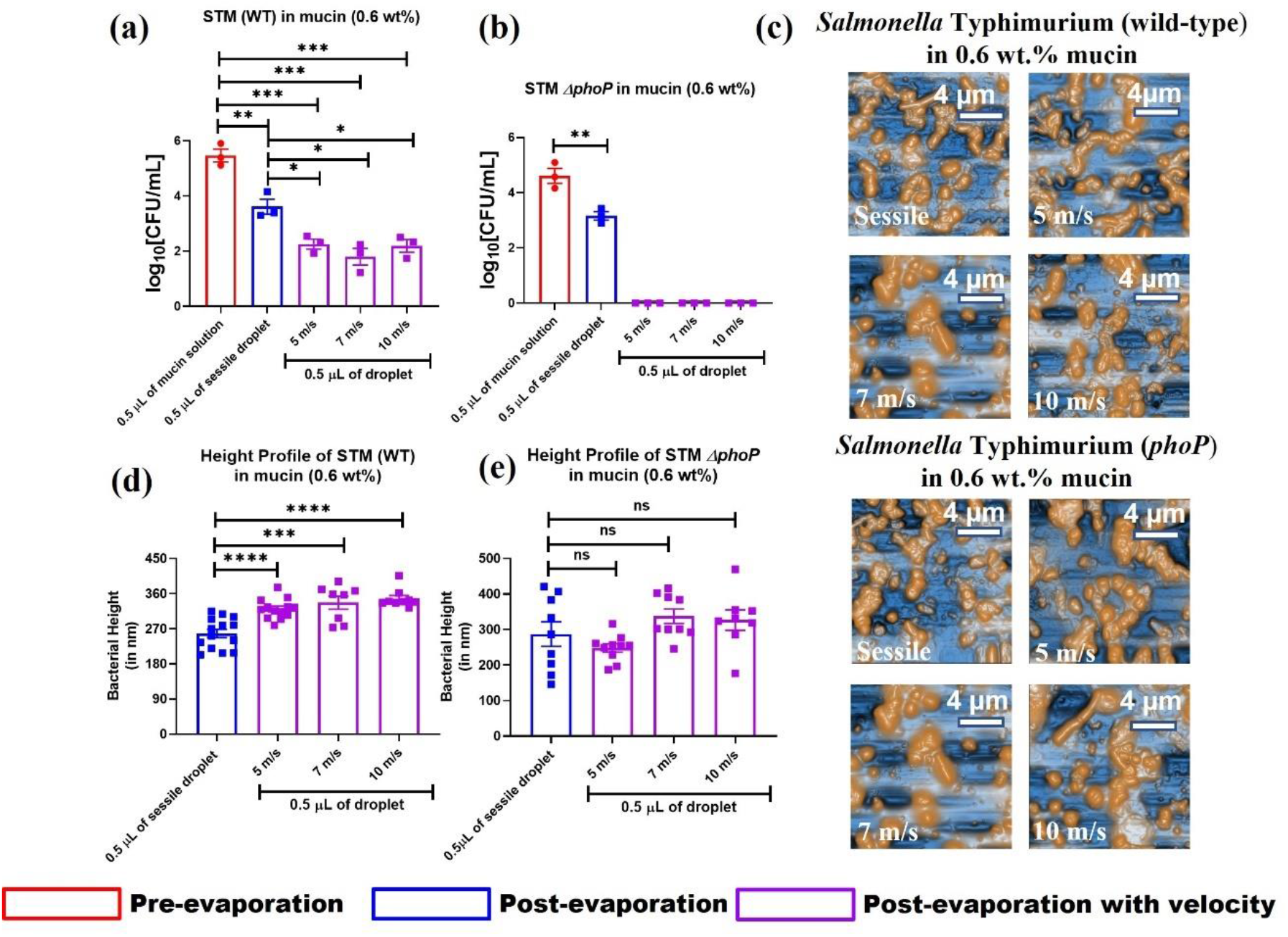
Deleting *phoP* reduced the *in vitro* viability of *Salmonella* Typhimurium in desiccated droplet of mucin (0.6 wt%) impacted on glass surface with velocity (5, 7, and 10 m/s). The *in vitro* viability of **(a)** STM (WT) and **(b)** *ΔphoP* log10 scale) recovered from dried sessile droplets of mucin (0.6 wt%) impacted solid glass surface with or without velocities (5, 7, and 10 m/s) **(N≥3)**. **(c)** AFM images of bacteria (STM WT and *ΔphoP*) laden droplets in mucin (0.6 wt%) on the glass surface. Height profile of **(d)** STM (WT) and **(e)** *ΔphoP* from the desiccated droplets of mucin (0.6 wt%) on the glass surface. Data are represented as mean ± SEM. ***(P)* *< 0.05, *(P)* **< 0.005, *(P)* ***< 0.0005, *(P)* ****< 0.0001, ns= non-significant, (Student’s t test-unpaired)**

The inability of STM *ΔphoP* survival in the stressed environment of velocity impacted desiccated droplets shows a novel role of *phoP* in assisting bacterial pathogens in tolerating mechanical stress. The deformation of the bacterial cell walls is an intrinsic aspect of impact velocity that affects the bacterial cell-cell interactions and surface topography. As illustrated in earlier sections, bacterial cell deformation depends on a few key parameters such as impact velocity, evaporative stress, and adhesion energy. In the case of smaller impact velocity (5 m/s) of droplets, the stretched time scales delayed the deformation of the bacterial cell phase due to lower receding velocity (Table 1) after initial contact with the substrate. Likewise, the increase in height profiles with an increase in the impact velocity is ascertained due to a decrease in the time scales of the receding phase, which results in the enhanced squeezing of bacterial cell walls. Interestingly, the height profile of STM *ΔphoP* (Fig. 6e) is strikingly different from the STM (WT) (Fig. 6d). The, STM *ΔphoP* exhibits an unresponsive height variation for an impact velocity of 5 m/s and the sessile drop (Fig. 6e). Similarly, a negligible increase is noted in STM *ΔphoP* even when the velocity is ramped up to 7 and 10 m/s. This indicates that the change in height profiles of *Salmonella* in velocity impacted desiccated droplets depends profoundly on *phoP* and, to a lesser extent, on mechanical stresses induced due to the impact.

In conclusion, the present study demonstrated that the investigation on bacteria-laden drop impact on surfaces is highly relevant from the disease spread perspective. This novel approach to droplet impact study opens a new dimension that brought out the effect of impact velocities on the survival possibilities of bacteria in the nutrient-rich and deficient medium. The spread of *Salmonella* from contaminated fomites or via aerosol is emerging as an important route of infections. Our study has discovered a few of the essential virulent genes of *Salmonella* that are required for their survival in desiccated droplets by bridging visible and microscopic perspectives. Even though the deformation of the bacterial cell wall is extensive at the higher impact velocity, the parameters such as receding velocity and adhesion energy also play an important role in controlling the deformation. An increase in bacterial cell height profile was noted for a higher impact Weber number due to high receding velocity occurring in smaller time scales. The enhanced adhesion energy in a nutrient-rich medium (0.1 and 0.6 wt.% mucin) provides cushioning effect compared to milli-Q bacterial cell walls to resist the mechanical stress induced due to the impact velocities. The better survival of wild-type *Salmonella* in the dried droplet of mucin (0.1 and 0.6 wt%) than in Milli-Q, glucose, and glycerol showed that the bacterial viability subjected to desiccation and mechanical stress induced by the impact velocity primarily depends upon the nutrient availability. The bacteria-laden droplet impact on the surface significantly reduces the viability of the bacteria during desiccation with the effect of additional mechanical stress. However, *Salmonella* survived in dried fomites efficiently proliferated in the macrophages. Our study revealed that, wild-type *Salmonella* accommodated with *phoP* in the desiccated droplets subjected to velocity-induced mechanical stress establishes a subsequent rise in infectivity in the macrophages.

## Methods

### Preparation of bacterial samples

Overnight grown 5 mL stationary-phase cultures of STM (WT) and *ΔphoP* (~10^8^ CFU) were pelleted down at 5000 rpm for 10 minutes. The cultures were washed twice with double autoclaved Milli-Q water and finally resuspended in 5 mL of neutral (Milli-Q), low-nutrient (glucose and glycerol), and nutrient-rich (LB broth, 0.1 and 0.6 wt% mucin) base solutions for further use.

### Side-view visualization

The LED light source (3W) is aligned in the direction of the Phantom Miro 110 camera to capture the drop impact phenomena for varying impact velocities. The images have been recorded at 10000 frames per second to measure droplet impact velocities and in-situ visualization of the droplet. The droplet impact velocities considered in this investigation (5, 7 and 10 m/s) were verified using this technique. The uncertainty of impact velocities is in the range of ±0.5 m/s.

### Interference Imaging

The bacteria-laden droplet spreading dynamics were captured using the reflection interference microscope during the impact on the glass surface for varying impact velocities. A laser light source of wavelength 640nm is in-line alignment with a beam splitter. An objective lens is placed at its focal length between the glass slide and the beam splitter for the fine focus of the light beam. The reflected beam from the beam-splitter passes through the bottom of the glass surface, and then the reflected light travels through the beam splitter to develop an interference fringe pattern, which is captured using high-speed imaging systems. The spreading dynamics of the drop impact have been recorded at 10000 frames per second simultaneously with the side-view imaging.

### Atomic Force Microscopy

A bio-Atomic Force Microscopy (AFM) (Park System, South Korea) integrated with an optical microscope with an X-Y flat scanner. The scanner is used in both contact and non-contact modes to acquire microscopic, scanned images, adhesion energy, and variation of force-distance spectroscopy data for different samples. The XEI, XEP software is integrated with the instrument used to operate, analyze, and store data to estimate various mechanical parameters. The technique of AFM measurement is based on laser beam deflection, where the laser beam is reflected from the rear side of the cantilever onto a position-sensitive detector. The position and force given to the sample are regulated using the instrumental software. ACTA cantilever with high stiffness and resonance frequency is used to get the topographical images of the sample. The contact mode AFM is carried out using CONTSCR (256px, scan rate 0.5 Hz) with less than 8 nm scanning radius. The stiffness of the cantilever is 0.2 N/m with a resonance frequency of 25kHz for acquiring the force indentation curve. Cantilever sensitivity and spring constant is calibrated before each experiment runs.

### Atomic Force Microscopy Analysis

The initial AFM data acquired from the bacterial sample is analyzed using XEI software. The force-indentation curves were processed for baseline correction to nullify the cantilever-bending to determine the tip-sample contact point and indentation force accurately. The roughness was calculated by averaging the RMS roughness of three independent 5μm^2^ areas.

### Assessment of the *in vitro* viability of the bacteria

0.5 μL of the bacterial samples, prepared in different base solutions, as mentioned earlier, were dispensed on a sterile glass slide with the help of a micropipette. The bacteria droplet is impacted at three different velocities on the glass substrate using a *PICO Pμlse* microdispensing system (*Nordson, USA*) by applying suitable fluid pressure to achieve the desired velocity (5,7 and 10 m/s). The experiments are conducted at room temperature 25°C and relative humidity of 45%. Five hours after the desiccation, 100 μL sterile PBS was added to the top of the slide to resuspend the dried bacterial droplets of different solutions. After resuspension, this 100 μL PBS was directly plated on *Salmonella-Shigella* (SS) agar. 100 μL from the overnight-grown culture and freshly prepared bacterial suspensions were plated to calculate the bacterial count in 0.5 μL of planktonic culture. Sixteen hours post-plating, the viable bacterial load was enumerated.

### Infection of RAW264.7 macrophages

Wild-type *Salmonella* Typhimurium was retrieved from the desiccated dried droplet of mucin as described earlier and used to infect RAW264.7 macrophage cells. The infected cells were incubated in a 5% CO_2_ incubator for 30 minutes at 37°C temperature, followed by washing with sterile PBS to clear the unattached bacteria. The cells were further incubated with 100 μg/ mL of gentamycin for an hour and 25 μg/ mL of gentamycin till the end of the experiment. The cells were lysed with 0.1% Triton-X100 at 3 hours and 18 hours. The lysates were serially diluted with PBS and plated on SS agar. To calculate the bacterial fold proliferation, the CFU was obtained from 18hr. was divided with the CFU from 3 hr. The CFU obtained from 3 hr. were further divided with the pre-inoculum CFU to calculate the percent phagocytosis of macrophages.

### Isolation of RNA from infected macrophages

The infected macrophage cells were treated with 1 mL of TRIZol solutions for lysis and kept in −80°C overnight. The lysates were further subjected to chloroform extraction, and to the isolated aqueous layer, an equal volume of isopropanol was added. The RNA was precipitated by centrifuging the mixture of aqueous layer and isopropanol at 14000 rpm for 20 minutes. The RNA pellet was washed with 70% ethanol and then resuspended in DEPC water. The RNA was treated with RNase-free DNase and converted into cDNA with the PrimeScript cDNA synthesis kit. The cDNA was subjected to RT-qPCR using SYBR green qRT-PCR kit to estimate the transcript-level expression of *sifA, ssaV*, and *phoP*.

### Statistical Analysis

Each experiment has been independently repeated several times, as indicated in the figure legends (n=technical replicates and N= biological replicates). GraphPad Prism 8.4.3 software was used to generate the graphs and perform the statistical tests. All statistics were performed in the manuscript by unpaired students t-test. The p values below 0.05 were considered significant in all the studies.

## Notes

### Competing Interest Statement

The authors have declared no competing interest.

