## Supplementary for "PHYSIOLOGICAL CHARACTERISTICS OF BACTERIAL DROPLETS INDICATE A CATASTROPHIC CONSEQUENCE WITH AN INCREASE IN IMPACT VELOCITY"

### Supplementary information

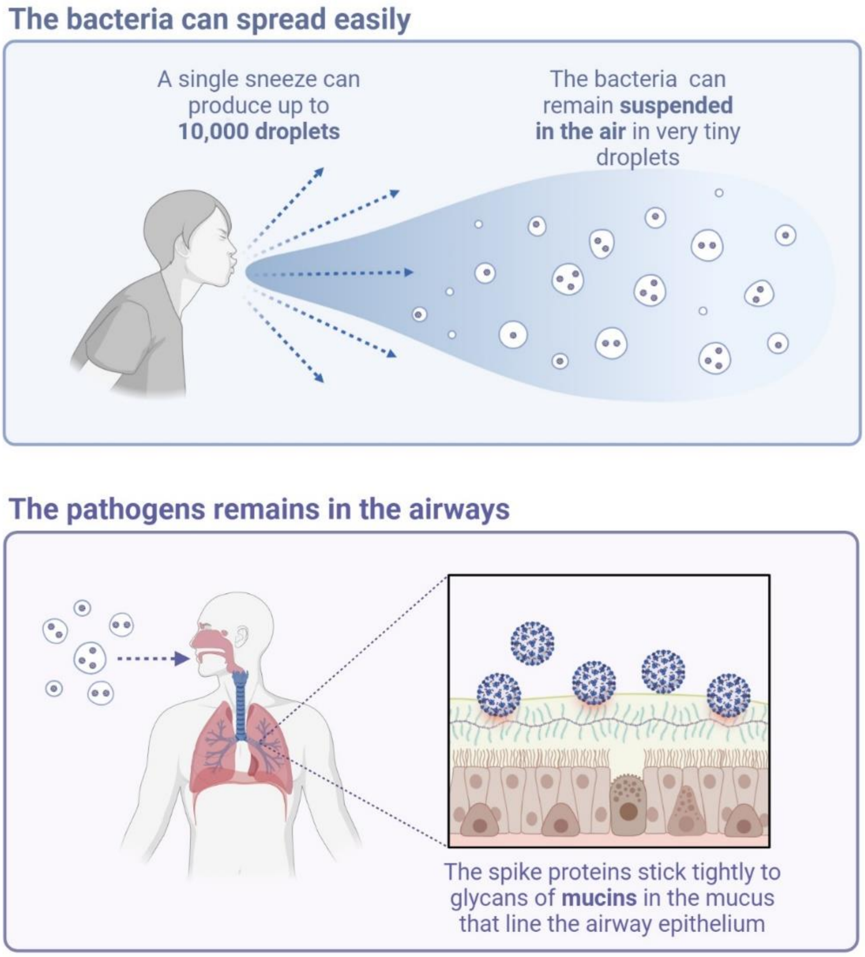


**Supplementary Figure 1.**

**Schematic diagram of fomites transmission.**
Schematic diagram indicating transmission of fomites from respiratory drops emitted during exhalation, coughing and sneezing. Diagram created with BioRender (www.biorender.com).

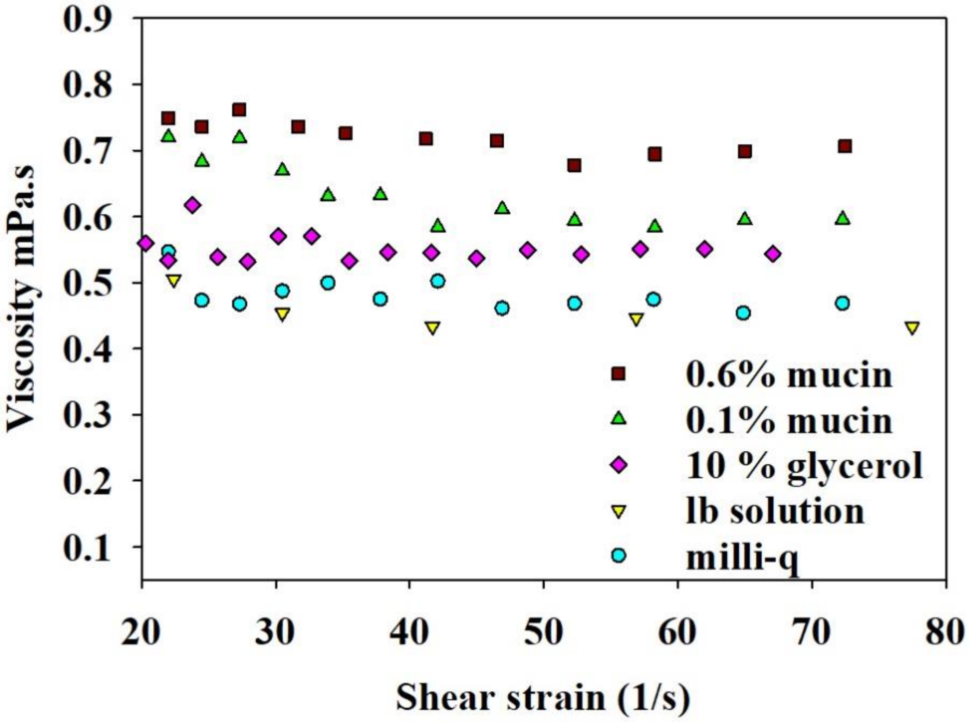


**Supplementary Figure 2.**

**Viscosity comparison of fluids shows a negligible variation.** Viscosity comparison for different samples milli-Q, 10% glycerol, LB solution, 0.6 wt% and 0.1 wt% mucin.

(a)

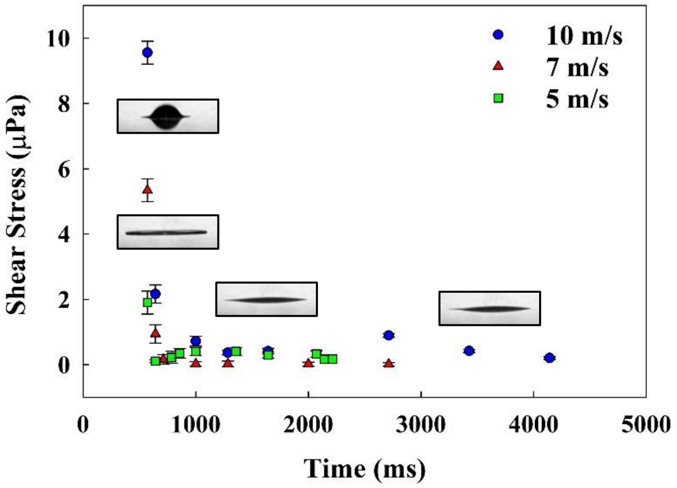


(b)

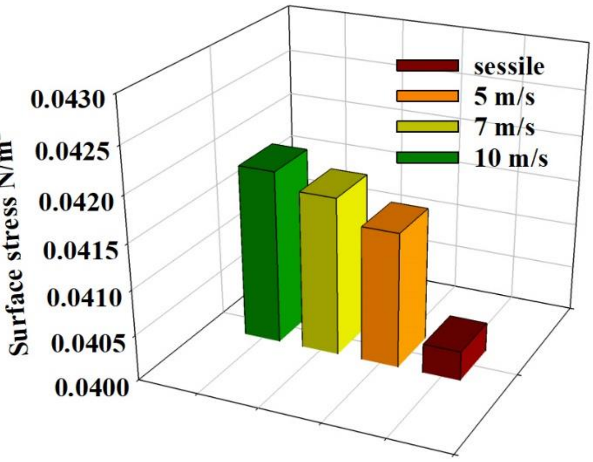


**Supplementary Figure 3.**
**Shear stress variation and surface stress experienced by AFM probe.**

The shear stress variation for the impacted sample on glass substrate and surface stress experienced by the AFM probe on four different impact conditions with milli-Q as the base medium.

**(a)**

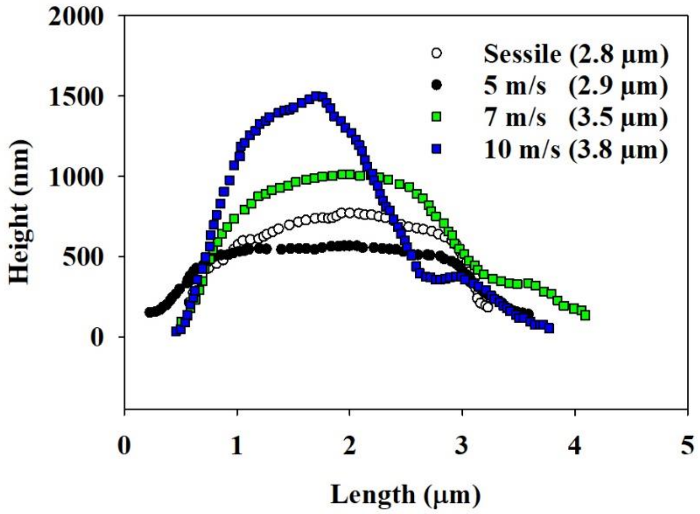


**(b)**

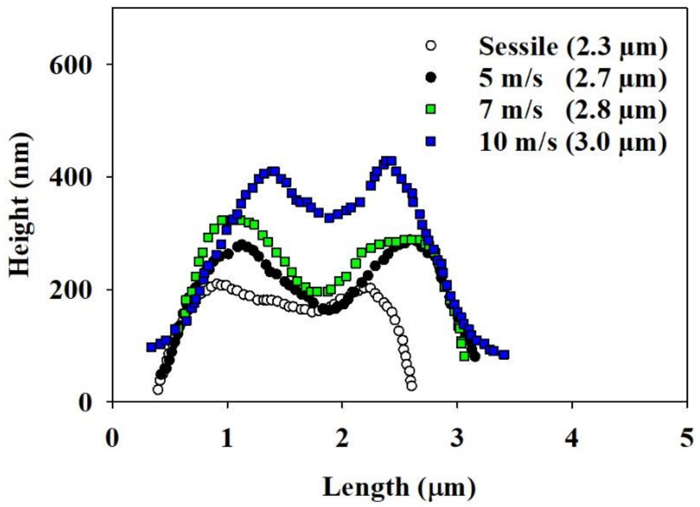


**Supplementary Figure 4.**
**Height and length profile of a random bacteria under different impact conditions.**

**(a)** The height profile of Salmonella Typhimurium in milli-Q subjected to different impact conditions. **(b)** The height profile of Salmonella Typhimurium in 0.6% mucin subjected to different impact conditions

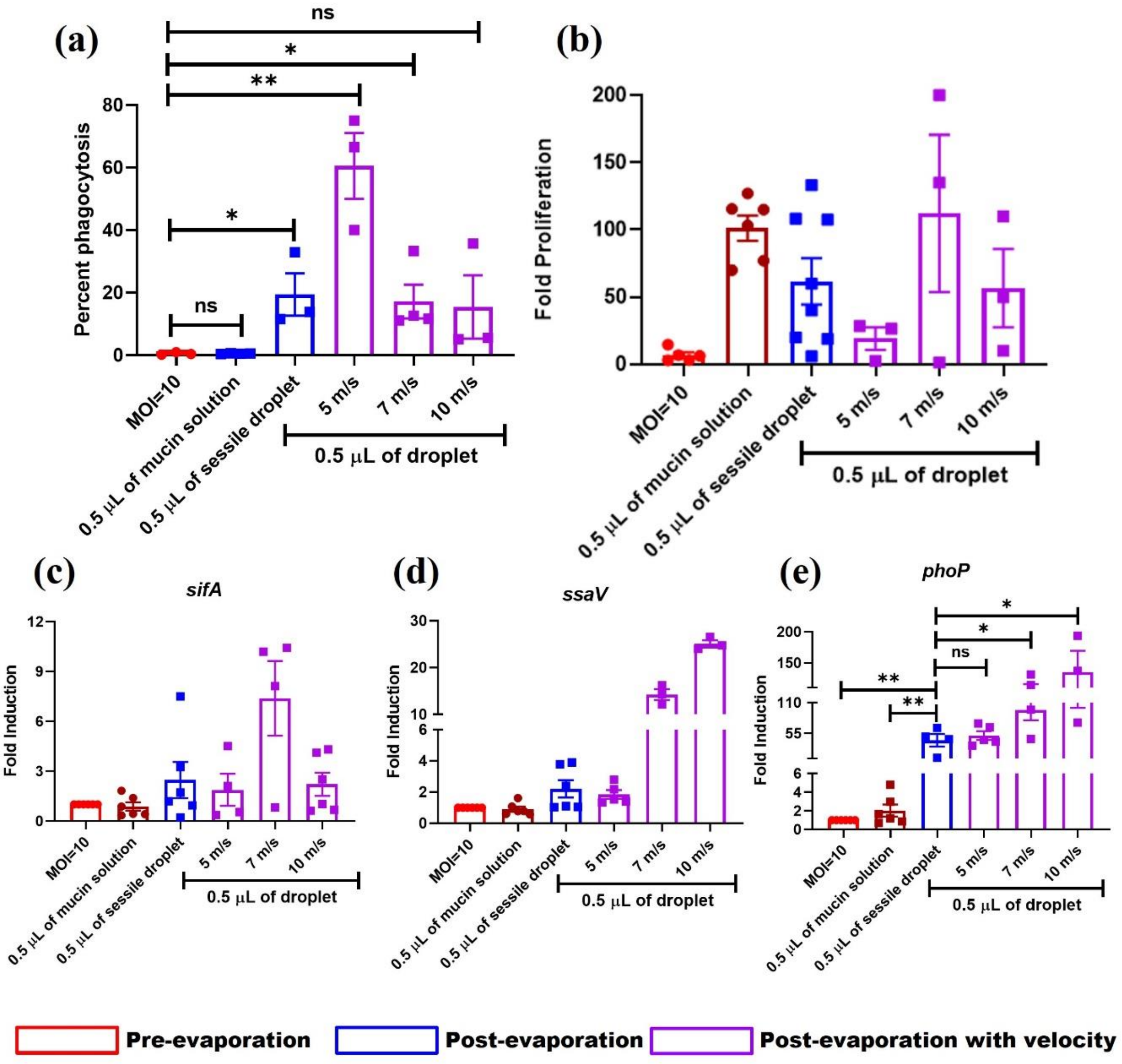


**Supplementary Figure 5.**
**The virulence of *Salmonella* Typhimurium retrieved from desiccated mucin droplet (0.6 wt%) impacted solid surface with or without velocities**

*Salmonella* Typhimurium recovered from desiccated mucin (0.6 wt%) droplet impacted solid glass surface with or without specific velocities (5, 7, and 10 m/s) were used to infect RAW264.7 cells to determine the **(a)** percent phagocytosis and **(b)** intracellular proliferation (n=3, N=2). Determining the transcript-level expression of **(c)** *sifA*, **(d)** *ssaV* and **(e)** *phoP* from intracellular *Salmonella* by RT-qPCR (n=3, N=2). Data are represented as mean ± SEM.

***(P)* *< 0.05, *(P)* **< 0.005, *(P)* ***< 0.0005, *(P)* ****< 0.0001, ns= non-significant, (Student's t test-unpaired)**
